# *OFF*-Target Effects of Mycophenolic Acid on BK Polyomavirus Replication in Primary Human Renal Proximal Tubular Epithelial Cells

**DOI:** 10.64898/2026.01.21.700791

**Authors:** Caroline A. Hillenbrand, Dorssa Akbari Bani, Océane M. Follonier, Zongsong Wu, Marion Wernli, Fabian H. Weissbach, Hans H. Hirsch

## Abstract

Tacrolimus (TAC) and mycophenolic acid (MPA) effectively reduce allograft rejection in kidney transplant recipients by inhibiting donor-specific lymphocyte activation and proliferation, respectively. However, this desired *on-target* effect is associated with uncontrolled BK polyomavirus (BKPyV)-replication and premature kidney allograft failure in 10%-20% of recipients. Besides impairing BKPyV-specific immunity, TAC also stimulates BKPyV-replication directly in renal proximal tubule epithelial cells (RPTECs) as an *off-target* effect. We now investigated *off-target* effects of MPA on BKPyV-replication in RPTECs. Following BKPyV exposure for 2 h, MPA inhibited BKPyV replication as shown by reduced supernatant BKPyV loads (IC_50_ 0.65 μM), number of infected cells, and viral protein expression at 72 hours post-infection (hpi). Notably, MPA inhibition of the viral large tumor antigen (LTag) was associated with the appearance of a truncated Tag of ∼17 kDa (truncTag-17). Adding guanosine (GUO) reversed MPA inhibition of BKPyV replication and viral protein expression, while truncTag-17 disappeared. Time course studies indicated that MPA inhibition was operative from 24 h before up to 24 h after BKPyV infection. GUO reversed MPA inhibition when administered from 24 h before to 24 h after BKPyV infection. Deep sequencing of passaged supernatant BKPyV genomes revealed more guanine-replacement (C→T or G→A) mutations for MPA compared to MPA/GUO (35% versus 10%, read threshold >0.5%). Thus, MPA can exert direct *off-target* effects on BKPyV replication in RPTECs linked to truncTag-17 expression, which can be partially reversed by exogenous GUO.

**Importance:** Reducing immunosuppression is currently recommended for kidney transplant recipients with BKPyV-DNAemia and nephropathy to improve virus-specific immune control, but increases the risk of T-cell and antibody-mediated rejection. Although TAC and MPA synergize in their *on-target* effects on lymphocytes, they differ in their *off-target* effects on RPTECs. Unlike TAC, MPA inhibits BKPyV replication in an uncompetitive dose-dependent manner. MPA inhibition can be partially reversed by exogenous GUO using the purine salvage pathway that is lacking in lymphocytes. MPA acts during the early viral replication phase that depends on LTag promoting cellular G1→S-phase progression. MPA-mediated GUO depletion not only reduces rapid cell proliferation similar to the well-known gastrointestinal or hematopoietic toxicities, but also slows viral replication, induces expression of a truncTag-17, and increases GC-mutation rates in progeny virus genomes. We discuss the relevance of these *off-target* characteristics of MPA in the optimized management of kidney transplant recipients with BKPyV-DNAemia and -nephropathy.

## Introduction

BK polyomavirus (BKPyV) was discovered in the 1970s in the urine of a kidney transplant (KT) recipient shedding morphologically altered “decoy cells” that harbored abundant polyomavirus-like particles in their enlarged nuclei ^1,2^. Although the clinical role of BKPyV in humans remained elusive for decades, BKPyV is linked today to three major complications occurring almost exclusively in immunocompromised hosts, namely BKPyV-associated nephropathy in 1%-15% after kidney transplantation ^3^, BKPyV-associated hemorrhagic cystitis in 5%-25% after allogeneic hematopoietic cell transplantation ^4^, and BKPyV-associated urothelial carcinoma in <0.1% of mostly kidney transplant recipients following chromosomal integration of the viral genome during prolonged high-level BKPyV replication ^5,6^. BKPyV-nephropathy is notable because of its sudden emergence in the late 1990s, after tacrolimus (TAC) and mycophenolic acid (MPA) replaced cyclosporine and azathioprine as standard maintenance immunosuppression ^7–10^. While TAC plus MPA combinations are associated with reduced rejection episodes during the first year post-transplant ^11^, poorly controlled BKPyV replication now contributes to premature kidney transplant failure in 10%–50% of affected KT recipients ^12–15^. Since no effective antivirals are available, current mitigation strategies recommend monthly screening for new-onset BKPyV-DNAemia and promptly reducing maintenance immunosuppression ^3^. Several factors have been associated with new-onset BKPyV-DNAemia and nephropathy in KT recipients including donor factors such as donor viruria, higher number of HLA-mismatches, and AB0-incompatibilty ^16–18^, recipient factors such as older age, male sex, low or undetectable BKPyV-specific cellular and humoral immunity ^19^ and insufficient coverage of all four BKPyV genotypes ^20–22^, as well as post-transplantation factors such as the use of TAC and MPA, high-dose glucocorticoid exposure ^10,23^, or non-use of mTOR inhibitors ^24^.

New-onset BKPyV-DNAemia and nephropathy in KT recipients have been largely viewed as an undesirable side-effect of the increased *on-target* immunosuppressive potency of TAC or MPA that reduced the allograft rejection rates ^19,25^. However, in larger randomized controlled studies reporting comparable immunologic potency as measured by rejection and graft failure, TAC and MPA combinations were associated with more BKPyV events compared to cyclosporine or mTOR-inhibitor combinations ^23,24,26–28^. While calcineurin inhibitors such as TAC and cyclosporin, and antiproliferative drugs such as mycophenolate, or mTOR-inhibitors sirolimus and everolimus synergize in their immunosuppressive *on*-*target* effects in lymphocytes, they can markedly differ in their *off-target* effects in non-immune cells. Thus, TAC stimulates BKPyV replication in primary human renal tubular epithelial cells (RPTECs), whereas the mTOR inhibitor sirolimus inhibits BKPyV replication through mechanisms requiring the intracellular binding protein FK-binding protein of 12 kDa known as FKBP-12 common to both, TAC and sirolimus ^29,30^. Unlike TAC, the calcineurin inhibitor cyclosporine has been reported to inhibit BKPyV replication through pathways requiring cyclophilin-A and NFATc3 ^30–32^.

The potential *off-target* effects of MPA are less well defined. However, MPA can affect rapidly dividing cells and may cause diarrhea and leukopenia as well-known *off-target* effects in gastrointestinal and hematopoietic cells, respectively. MPA is a reversible uncompetitive inhibitor of the inosine monophosphate dehydrogenase (IMPDH), the rate-limiting enzyme of the *de novo* purine biosynthesis. IMDPH converts inosine monophosphate (IMP) to xanthosine monophosphate (XMP) ^33^. Since lymphocytes lack the salvage pathway of purine synthesis and solely rely on the *de novo* pathway, MPA acts as an antiproliferative immunosuppressive drug. Here, we investigated potential *off-target* effects of MPA on BKPyV replication in primary human RPTECs.

## Results

### MPA treatment inhibits BKPyV progeny release in primary human RPTECs

To investigate the effects of MPA on BKPyV replication, we incubated primary human RPTECs for 2 h with BKPyV-Dunlop at 24 h after seeding (**Fig. 1A**). After washing to remove unbound virions, the cells were cultured in the presence of increasing concentrations of MPA or solvent (DMSO). At 72 h post-infection (hpi), cell culture supernatants were incubated with DNase-I to remove non-encapsidated viral genomes prior to total nucleic acid extraction and analyzed for BKPyV-DNA loads by quantitative nucleic acid testing (QNAT) (see Materials & Methods)^34^. The results showed that MPA treatment decreased the BKPyV loads in the supernatants in a dose-dependent manner (**Fig. 1B**). Compared to the solvent control taken as a 100% reference, MPA concentrations of >1 µM were associated with significantly reduced viral loads (**Fig. 1C**). In the curve-fitting model, the inhibitory concentration 50 (IC_50_) was estimated as 0.65 µM of MPA (**Fig. 1D**). However, the inhibition leveled off at higher MPA concentrations and the IC_90_ value could not be calculated. Thus, the MPA treatment reduced BKPyV supernatant loads consistent with an uncompetitive inhibition pattern that had been reported for the effect of MPA on the IMPDH activity ^35,36^. We concluded that MPA reduced the release of viral progeny into cell culture supernatants in a dose-dependent, but uncompetitive manner with an estimated IC_50_ of 0.65 µM.

**Fig. 1:**
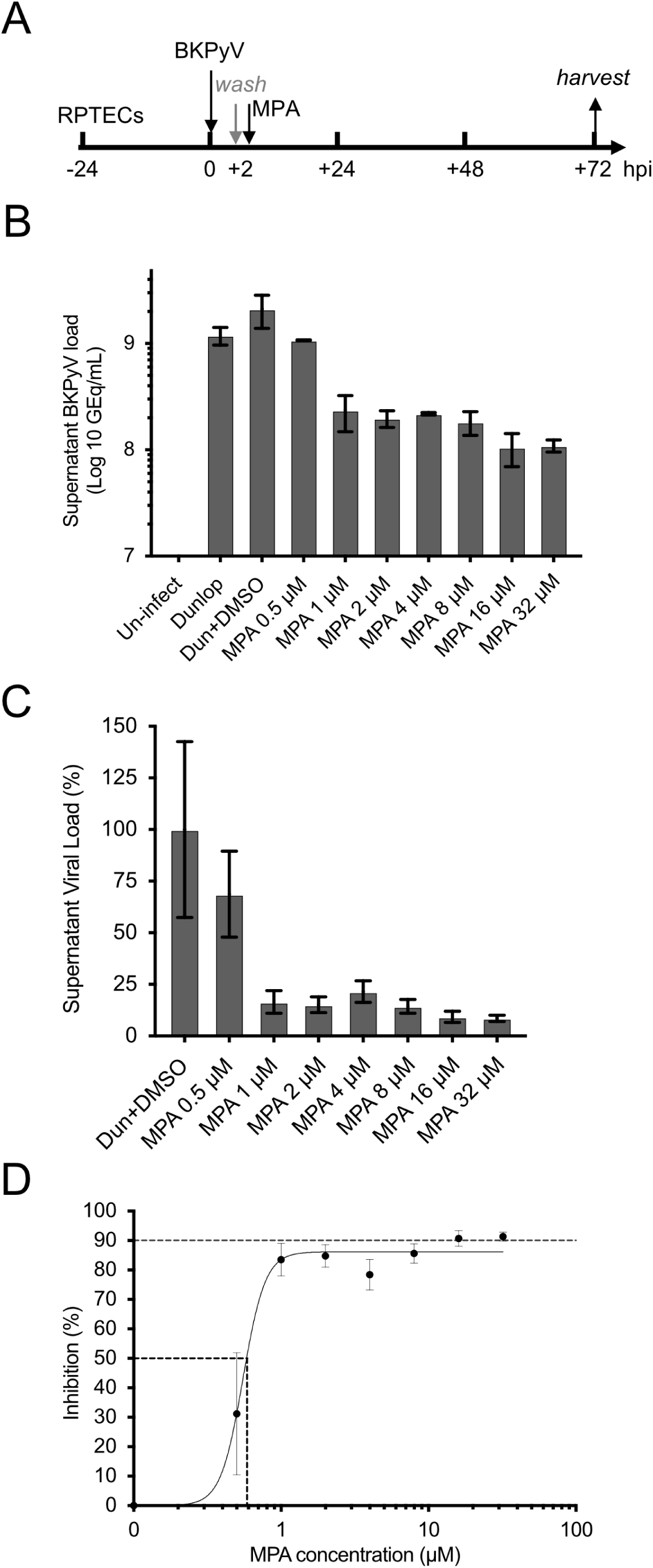
MPA inhibits BKPyV replication in primary human RPTECs. The conditions of cell seeding, BKPyV infection, MPA treatment and determination of supernatant viral loads are detailed in Material & Methods. A. Time line of experimental procedures including cell seeding, virus exposure, MPA addition and harvest for analysis. B. Determination of BKPyV DNA loads in cell culture supernatants at 72 hpi for the indicated conditions. C. Percentage of supernatant viral loads relative to the solvent control from panel B. D. IC_50_ estimates using a curve-fitting model based on percent viral loads in panel C.

### MPA treatment reduces the number of BKPyV-infected RPTECs

To investigate whether the reduced supernatant BKPyV loads resulted from impaired virus infection or production, we performed immunofluorescence microscopy (Materials & Methods). RPTECs were seeded on coverslips 24 h before BKPyV infection and treated with increasing MPA concentrations as outlined above (**Fig. 1A**). At 72 hpi, the adherent cells were fixed and immunofluorescence microscopy was performed for the viral large T-antigen (LTag), the major capsid protein Vp1, agnoprotein as well as cell nuclei (**Fig. 2A**)^37^. Compared to the solvent control (0 μM), increasing MPA concentrations were associated with decreasing numbers of BKPyV-positive RPTECs. At MPA 0.625 μM, the number of LTag-positive or Vp1-positive cells were reduced by approximately one half in line with the calculated IC_50_ found for the supernatant BKPyV-DNA loads. With increasing MPA concentration, the total number of cells were also reduced. The results indicated that higher MPA concentrations not only reduced BKPyV-supernatant loads at 72 hpi, but also the number of BKPyV-infected and uninfected cells.

**Fig. 2:**
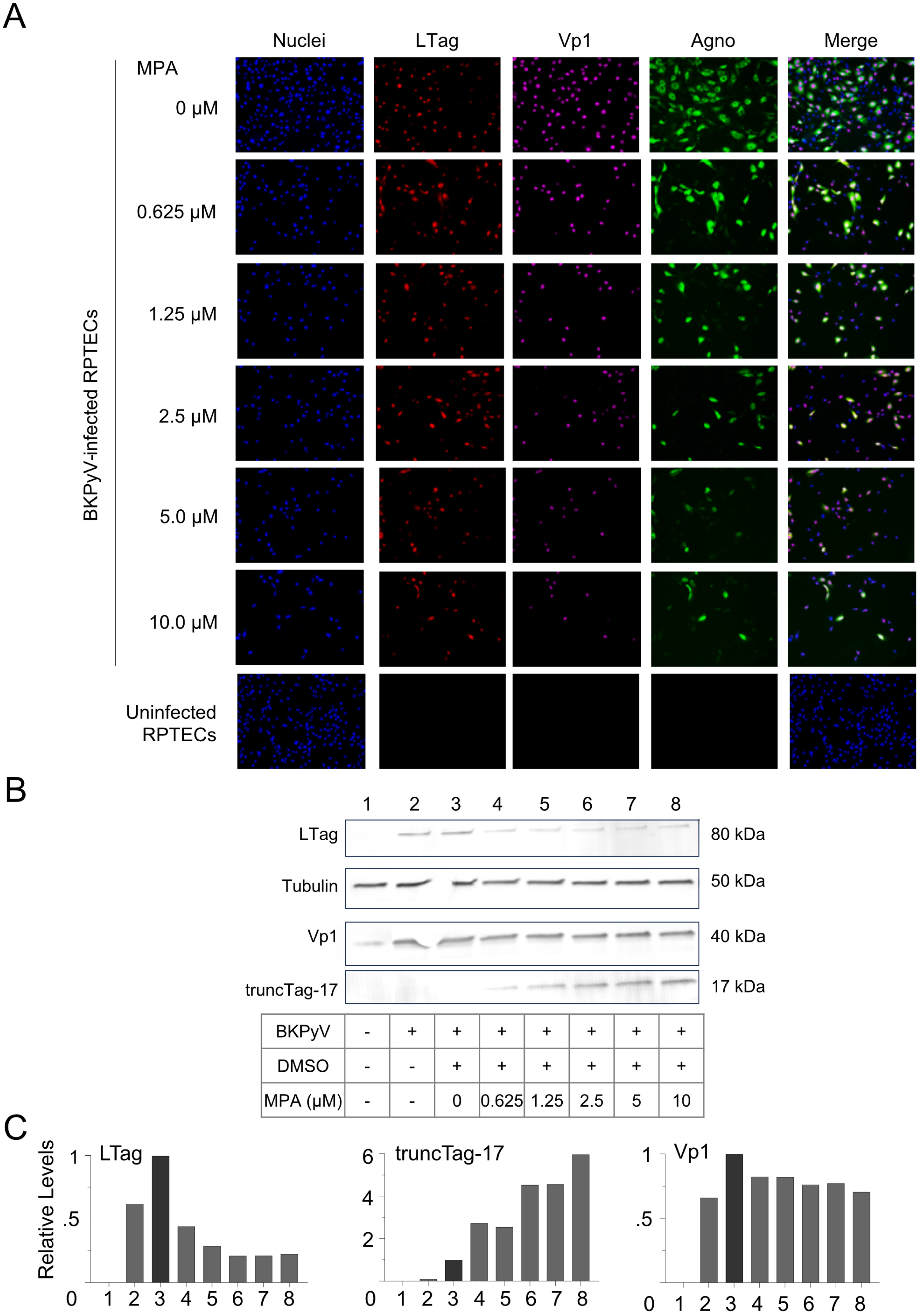
MPA reduces the number of BKPyV-positive cells and promotes truncTag-17 expression. RPTECs were seeded, infected with BKPyV and treated with the indicated concentrations of MPA at 2 hpi, as outlined in Fig. 1A (see Materials & Methods). A. Immunofluorescence microscopy of RPTECs at 72 hpi. B. Immunoblot analysis of cell lysates harvested at 72 hpi. C. Relative levels of LTag, Vp1, and truncTag-17, normalized to tubulin.

### MPA treatment reduces LTag levels and leads to the appearance of truncated Tag of 17kDa

To examine the impact of increasing MPA concentrations on viral protein expression more directly, we performed immunoblots from adherent cells harvested at 72 hpi (**Fig. 2B; Supplementary Fig. 1**). The normalized protein bands corresponding to LTag and Vp1 declined as the MPA concentrations increased. Interestingly, the decrease in LTag after MPA addition was associated with the appearance of a truncated antigen of approximately 17 kDa (truncTag-17) also detected by the anti-LTag antibody (**Fig. 2B; Supplementary Fig. 1**). The truncTag-17 increased in parallel to the MPA concentrations, but was virtually undetectable in the DMSO solvent control. Quantification of the tubulin-normalized bands showed that LTag levels decreased 2.5- to 5-fold to approximately 20%-40%, whereas Vp1 expression only decreased by 1.5-fold to 60%-70% (**Fig. 2C; Supplementary Fig.1**). In contrast, the truncTag-17 increased by approximately 5-fold over the background. We concluded that MPA treatment inhibited BKPyV replication as demonstrated by reduced supernatant viral loads, decreased viral protein expression, whereby LTag expression was more reduced than Vp1 and coupled to the appearance of a novel truncTag-17.

To address the inhibition of host cell proliferation at higher MPA concentrations, we noted that Tylden et al. ^38^ had reported more pronounced cytostatic effects for the antiviral brincidofovir, a lipid conjugate of cidofovir, when cells were seeded at lower density. Following improved cell adherence on coated cover slips after seeding, uninfected and BKPyV-infected RPTECs reached confluency at 72 hpi when cultured in the absence of MPA (**Fig. 3A**). At MPA concentrations of 0.5 μM and 1 μM, the BKPyV-infected cell cultures were slightly less confluent at 72 hpi. At MPA concentrations of 2 μM and more, the confluency of the RPTEC cultures was clearly reduced. Immunofluorescence staining revealed that at MPA concentrations of 2 μM and more, the total number and the number of LTag-expressing cells were decreased (**Fig. 3B**). Notably, the cytoplasmic agnoprotein and the nuclear Vp1 capsid protein were detected in a substantial number of cells. We concluded that MPA addition at 2 hpi appeared to affect the number of infected cells as well as the total number of cells detectable at 72 hpi, whereby a residual population remained refractory to MPA inhibition.

**Fig. 3:**
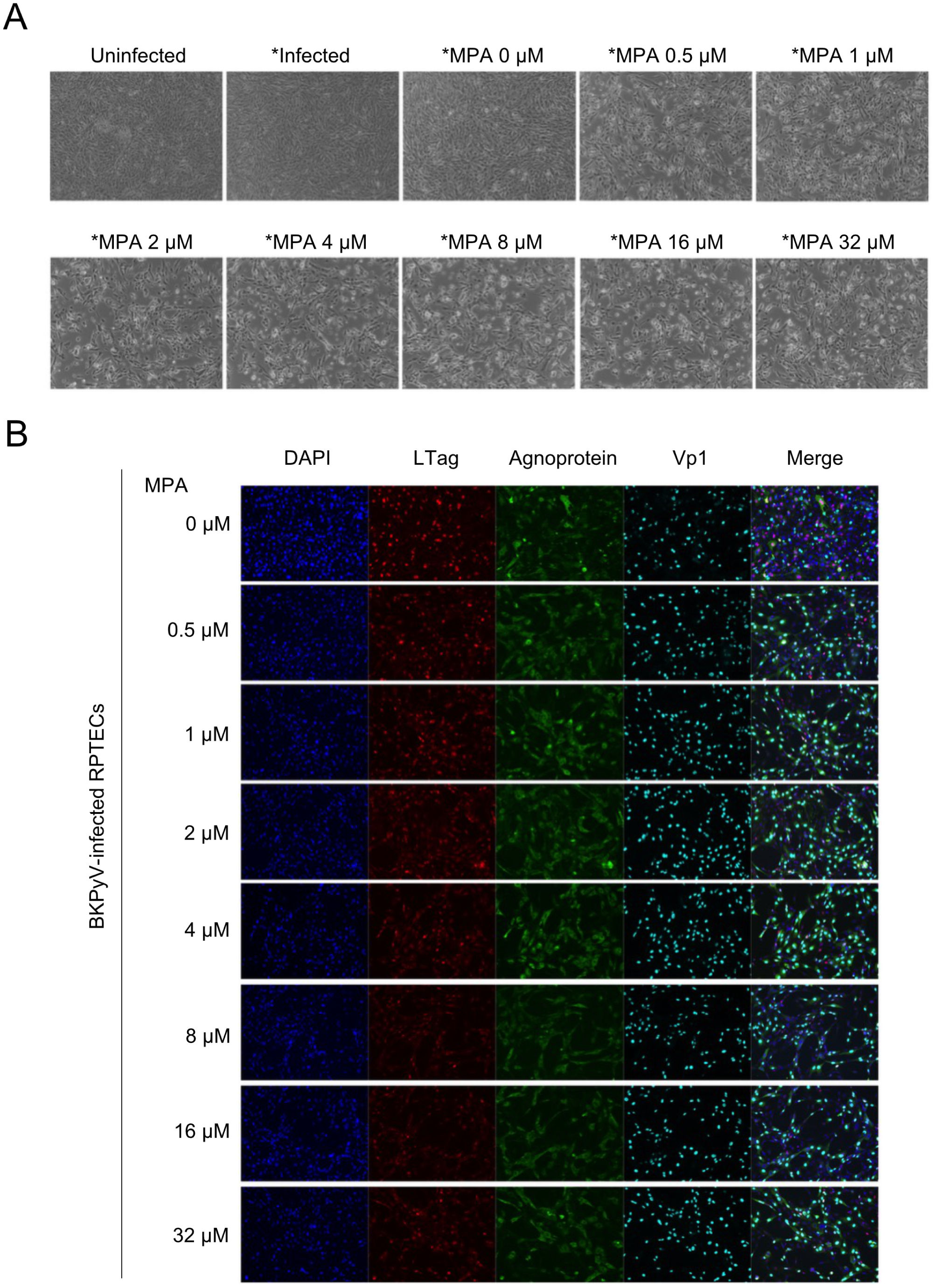
MPA reduces confluency and number of BKPyV-infected cells. RPTECs were seeded and treated with the indicated concentrations of MPA (0 - 32 μM) at 2 hpi, as outlined in Fig. 1A (see text and Materials & Methods). A. Confluency of RPTECs at 72 hpi by phase contrast microscopy. B. Immunofluorescence microscopy of RPTECs at 72 hpi treated with the indicated concentrations of MPA.suppl

### MPA inhibition of BKPyV-replication in RPTECs is reversed by exogenous guanosine

To investigate the effects of MPA on the dynamics of RPTEC proliferation, we measured the impedance of adherent cell monolayers using the xCELLigence system ^38,39^, which allows for real-time monitoring of the cell index (**Fig. 4A**). The cells were seeded and infected with BKPyV after 24 h, and followed until 72 hpi for the indicated conditions. In the presence of medium or solvent control, the cells proliferated and nearly doubled their cell index. In the presence of MPA, the cell index was not increasing but rather declined slowly. To investigate if the MPA inhibition could be reversed by exogenous guanosine (GUO), BKPyV-infected RPTECs were treated with MPA 2.5 μM at 2 hpi followed by adding the indicated concentrations of GUO at 6hpi (**Fig 4B**). The results showed that despite this four-hour delay, exogenous GUO largely reversed the MPA inhibition of RPTEC proliferation.

**Fig. 4:**
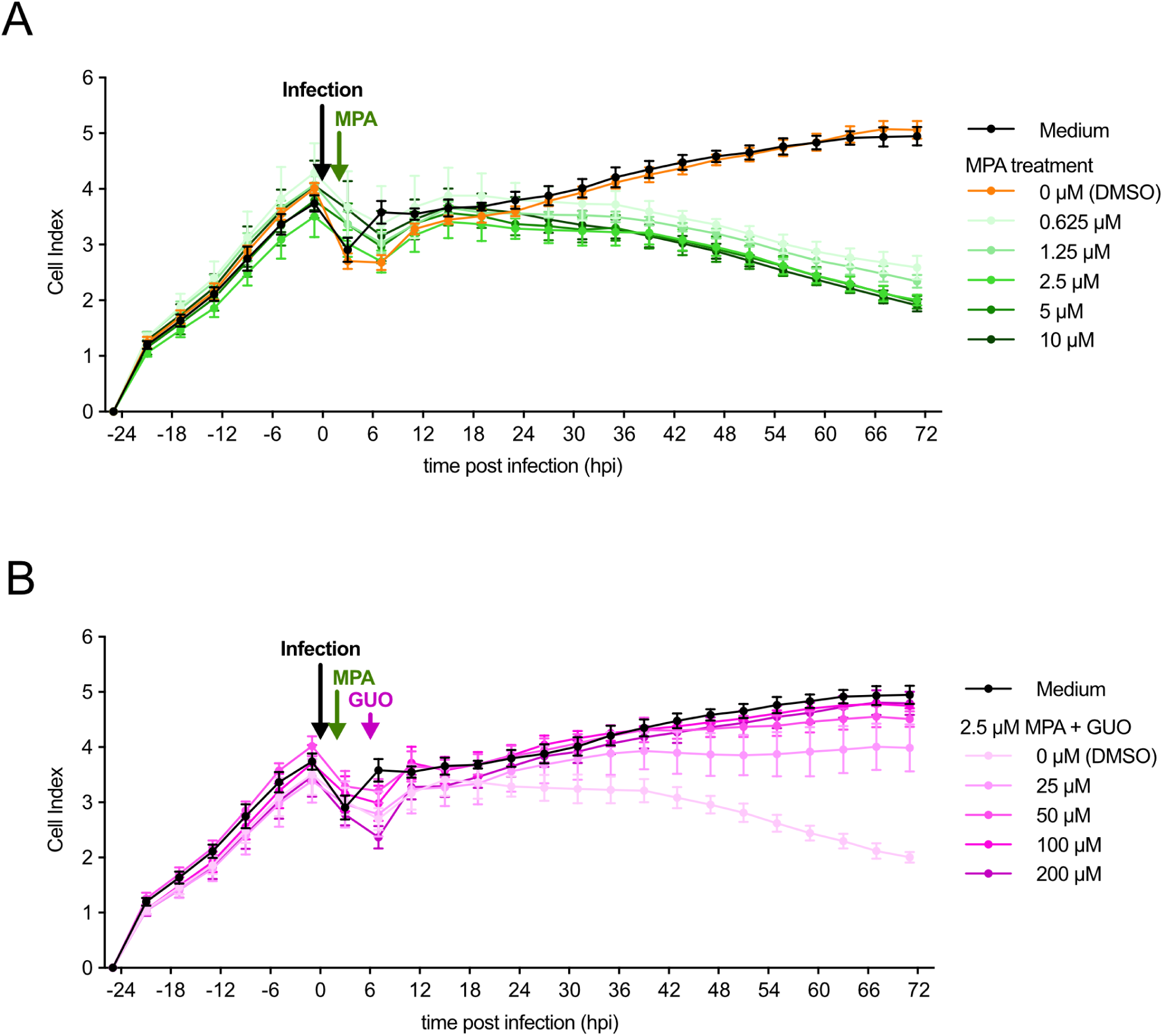
Proliferation of BKPyV-infected primary human RPTECs is impaired by MPA, but can be reversed by guanosine (GUO) supplementation. RPTECs were seeded, infected with BKPyV, treated with the indicated concentrations of MPA at 2hpi and GUO at 6 hpi (see text and Materials & Methods). Cell indices were monitored in real-time by impedance measurements of the 96-well xCELLigence system A. MPA inhibits RPTEC proliferation as measured by real-time impedance B. MPA inhibition of RPTECs can be reversed by exogenous GUO.

To investigate whether exogenous GUO could also reverse the MPA inhibition of BKPyV replication, we added increasing concentrations of MPA at 2 hpi followed by 200 μM GUO at 6 hpi. Quantification of the supernatant viral loads at 72 hpi showed that exogeneous GUO reversed the MPA inhibition from 0.625 up to 5 μM, (**Fig. 5A**). Interestingly, the supernatant BKPyV loads in the presence of GUO were higher than in the solvent control. At 10 μM MPA, however, the supernatant BKPyV loads were approximately 20% lower than the control suggesting that GUO reversal faded at these higher MPA concentrations.

**Fig. 5:**
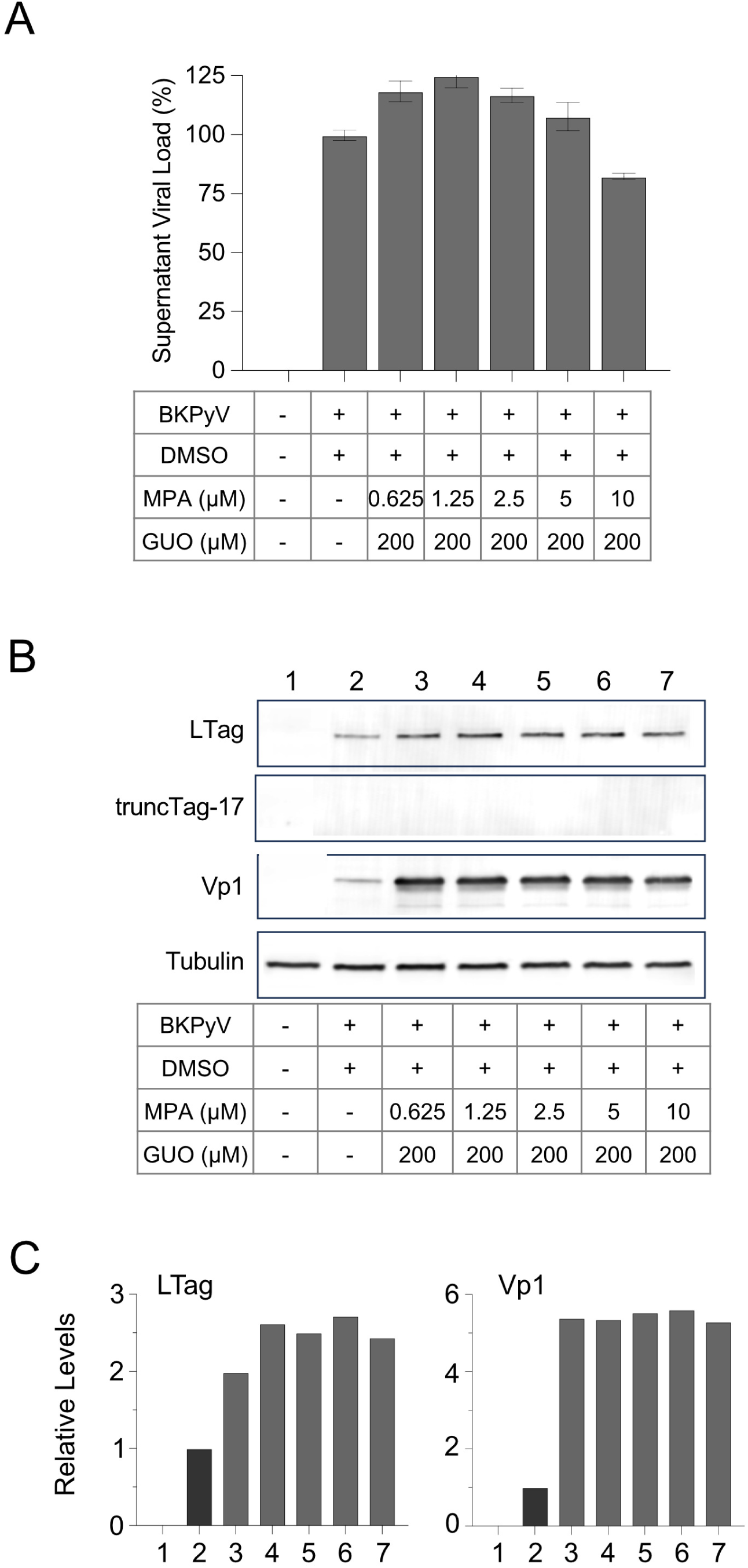
Exogenous GUO reverses MPA inhibition of BKPyV replication in primary human RPTECs. RPTECs were seeded, infected with BKPyV, treated with the indicated concentrations ofMPA at 2 hpi and 200 μM GUO at 6 hpi. A. Supernatant BKPyV-loads were determined by QNAT after DNase-I digestion and total nucleic extraction at 72 hpi as outlined in Fig 1. B. Immunoblot analysis of cell lysates at 72 hpi. C. Relative levels of LTag and Vp1, normalized to tubulin.

Immunoblots of cell extracts harvested at 72 hpi demonstrated that exogenous GUO reversed the inhibitory MPA effect on LTag and Vp1 levels, which were increased compared to the solvent control, whereas the truncTag-17 was not detectable under these conditions (**Fig. 5B; Supplementary Fig. 2**). Indeed, LTag and Vp1 showed a 3- to 5-fold and 6-fold increase, respectively, compared to the solvent control (**Fig. 5C**). We concluded that exogenous GUO addition could reverse MPA-inhibition of BKPyV replication even when added 4 hours later.

To examine the reversal potential of GUO at higher MPA concentrations, we compared the real-time proliferation of RPTECs seeded at −24 hpi, followed by addition of 8 μM MPA added at 2 hpi and increasing GUO concentrations added at 6 hpi (**Supplementary Fig. 3A**). The data showed a GUO concentration-dependent recovery of the cell index. Quantification of the supernatant viral loads at 72 hpi revealed a stepwise increase with increasing GUO concentrations, whereby the addition of 200 μM GUO at 6 hpi largely reversed the inhibition of BKPyV replication of MPA 8 μM given at 2 hpi (**Supplementary Fig. 3B**). Taken together, the data confirmed that increasing concentrations of exogenous GUO could largely reverse MPA-mediated inhibition of BKPyV replication and RPTEC proliferation even when added after a four-hour delay.

### The early phase of BKPyV replication is critical for GUO reversal of MPA inhibition

To investigate whether there is a limited time window for the GUO reversal of MPA-inhibition, RPTECs were treated with 2.5 μM of MPA at 2 hpi, and 200 μM of GUO was added at the indicated times before, at, or after BKPyV infection. Brightfield microscopy at 72 hpi showed the reduced confluency of the RPTECs when only MPA was added at 2 hpi. In contrast, dense confluency was seen when adding GUO from −24 hpi until +36 hpi, which only slightly decreased when adding GUO as late as 48 hpi (**Fig. 6A**). When analyzing DNase-protected supernatant BKPyV-loads at 72 hpi, GUO supplementation was able to restore viral replication when added between −24 hpi, 0 hpi, +2 hpi and +6 hpi, and only slightly reduced supernatant BKPyV load were seen when GUO was added at +12 hpi. However, later supplementation of GUO at +24 hpi, +36 hpi, and +48 hpi was associated with a stepwise reduction in DNase-protected supernatant BKPyV-loads towards approximately 20% of the levels seen at +6 hpi (**Fig. 6B**). GUO-reversal of MPA-inhibition was similarly seen in DNase-untreated supernatant BKPyV loads.

**Fig. 6:**
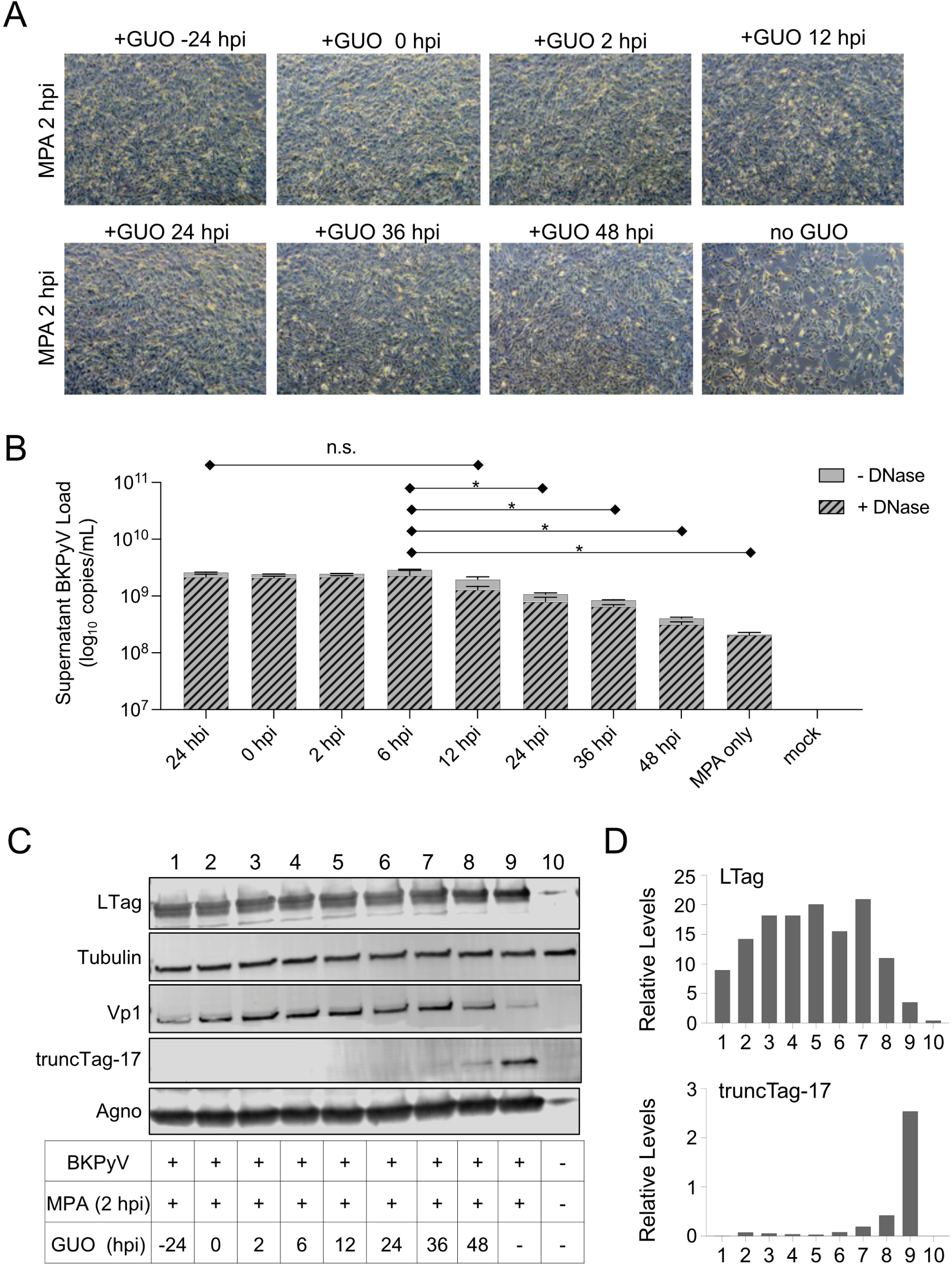
Time course of GUO supplementation to reverse MPA inhibition of BKPyV replication in primary human RPTECs. RPTECs were seeded, infected with BKPyV, treated with the indicated concentrations of MPA at 2 hpi and 200 μM GUO was added at the indicated time points. A. Confluency of RPTECs at 72 hpi by phase contrast microscopy. B. Effect of GUO supplementation at the indicated time points on supernatant BKPyV-loads. BKPyV-loads were determined by QNAT without (-DNase) or with (+DNase) digestion by DNase-I prior to total nucleic extraction at 72 hpi (DNase-treated; * p < 0.05, one-way Anova) C. Viral protein expression by immunoblot analysis at 72 hpi. Cell lysates were prepared and analyzed by immunoblots using antibodies against LTag, tubulin, Vp1, truncTag-17, and agnoprotein. D. Relative levels of LTag and truncTag-17, normalized to tubulin.

Immunoblot analysis at 72 hpi showed that GUO supplementation until +36 hpi was effective in maintaining BKPyV LTag levels (**Fig. 6C; Supplementary Fig. 4**). At this timepoint, the truncTag-17 band started to appear and increased in intensity until GUO addition at 48 hpi, but the highest levels of truncTag-17 were seen for MPA 2.5 μM without GUO supplementation. The late viral gene region-encoded Vp1 and agnoprotein were affected to a lesser extent. Quantification of the immunoblot results showed that the relative LTag levels were slightly higher when GUO was added after the washing steps to remove BKPyV as compared to earlier time points (**Fig. 6D**). We concluded that GUO was able to counteract MPA-inhibition of BKPyV replication, if the supplementation occurred before +24 hpi, i.e., before LTag induced the host cell changes preceding high-level viral genome replication at 36 hpi. Moreover, truncTag-17 was confirmed as a marker of MPA-induced GUO depletion and was inversely related to LTag levels.

### MPA acts during the early BKPyV replication phase in RPTECs

To investigate whether there is a limited time window for the MPA-inhibition, 2.5 μM MPA was added at the specified times before and after BKPyV infection (**Fig. 7**). Brightfield microscopy at 72 hpi showed that MPA was associated with significantly reduced confluence when added at −24 hpi, 0 hpi, and +2 hpi (**Fig. 7A**). This effect faded when MPA was added at 12 hpi, and nearly or fully confluent RPTEC monolayers were seen at 72 hpi, when adding MPA at 24 hpi or later, respectively. The supernatant BKPyV loads at 72 hpi were reduced by MPA addition until 12 hpi and showed a step-wise increase at the later time points 48 hpi (**Fig. 7B**). Immunoblot analysis of RPTECs at 72 hpi showed that the viral proteins LTag, Vp1, and agnoprotein increased with increasing delay of MPA addition (**Fig. 7C; Supplementary Fig. 5**). Conversely, truncTag-17 showed an initial increase before returning to background levels. Quantification of the immunoblot results showed that LTag levels increased gradually with increasing delay of MPA addition, reaching the highest levels at +48 hpi, comparable to the GUO rescue of MPA added at 2 hpi (**Fig. 7D**). We concluded that the inhibitory effect of MPA treatment was strongest when added within 12 hours of BKPyV infection. Thus, the time window of MPA-inhibition matched the ability of exogenous GUO to rescue BKPyV replication.

**Fig. 7:**
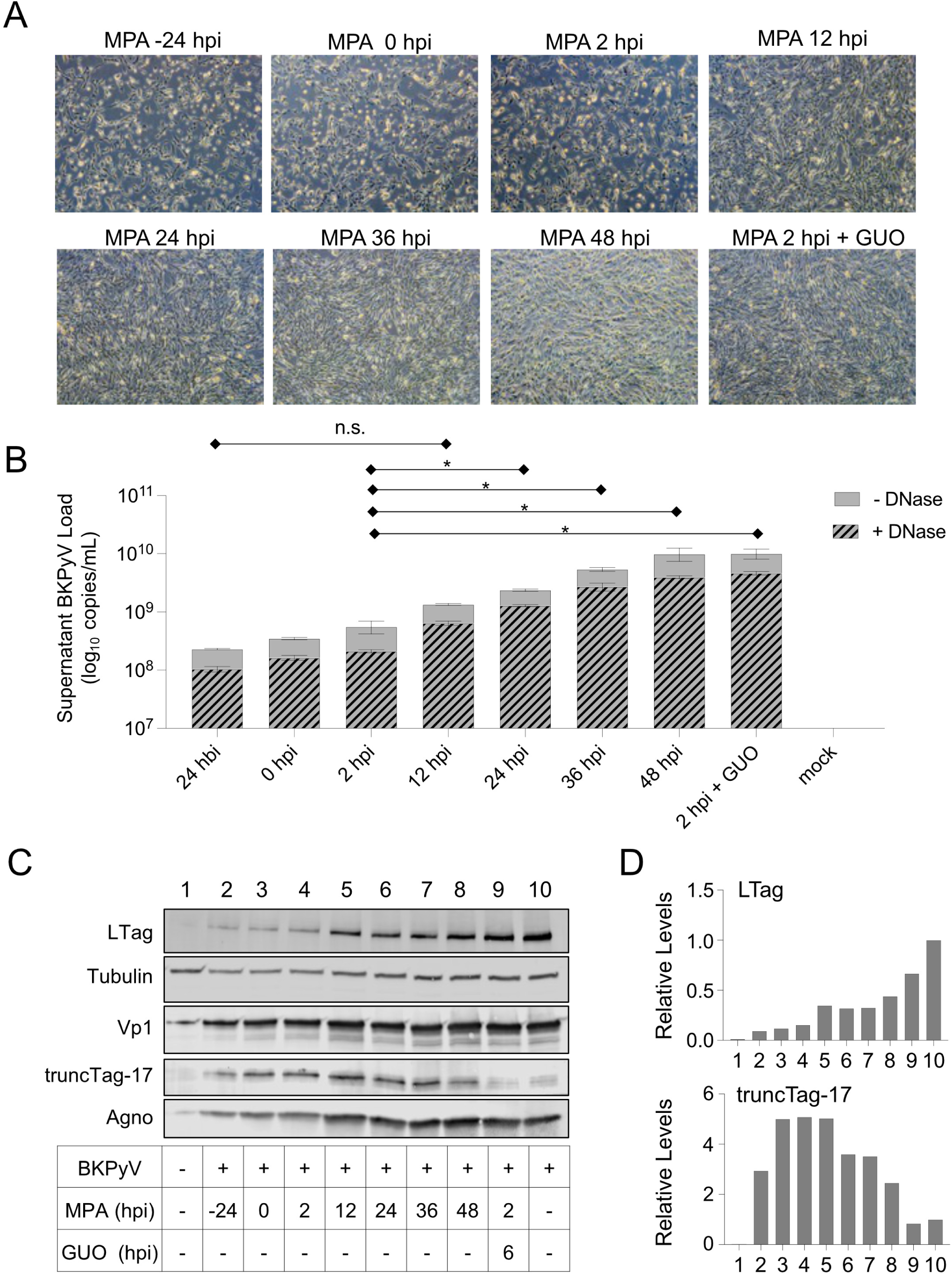
Time course of MPA addition to inhibit BKPyV replication in primary human RPTECs. RPTECs were seeded, infected with BKPyV, treated with the 2.5 μM of MPA at the indicated time points, or at 2 hpi followed by addition of 200 μM GUO at 6 hpi. A. Confluency of RPTECs at 72 hpi by phase contrast microscopy. B. Effect of MPA addition at the indicated time points on supernatant BKPyV-loads. BKPyV-loads were determined by QNAT without (-DNase) or with (+DNase) digestion by DNase-I prior to total nucleic extraction at 72 hpi (DNAse-treated; * p < 0.05, one-way Anova). C. Viral protein expression by immunoblot analysis at 72 hpi. Cell lysates were prepared and analyzed by immunoblots using antibodies against LTag, tubulin, Vp1, truncTag-17, and agnoprotein. D. Relative levels of LTag and truncTag-17, normalized to tubulin.

### MPA is associated with a higher rate of BKPyV-genome mutations

To explore whether MPA treatment-induced shortage of GUO was associated with an increase in viral genome mutations, we analyzed BKPyV genomes in the 72 hpi supernatants by next generation sequencing (see Materials & Methods). To this end, duplicate cultures of RPTECs were infected with BKPyV and either treated with 2.5 μM MPA at 2 hpi (MPA) or treated with 2.5 μM MPA at 2 hpi followed by the addition of 200 μM GUO at 6 hpi (MPA/GUO). The DNase-protected supernatant BKPyV loads at 72 hpi were determined for both conditions, adjusted to equal viral loads and used for two sequential rounds of the same infection conditions with fresh RPTECs. We observed that the supernatant BKPyV loads decreased for the MPA condition after each of three passages, but increased for the MPA/GUO condition. From the supernatants of passage 3, the total nucleic acids were extracted after DNase treatment and submitted to next generation sequencing of 10 gel-purified 1000 bp-amplicons covering the entire BKPyV genome (**Fig. 8**; **Supplementary Table 1**)^34^. Overall, more mutations were observed for BKPyV genomes under the MPA condition compared to the MPA/GUO condition. In fact, a greater proportion of guanosine-replacement mutations (C→T or G→A) was observed in two independent replicates of the MPA condition (40% and 36%) compared to 14% and 13% in the MPA/GUO condition (**Fig. 8A**). The mutations were detected in all viral genome regions and included non-synonymous changes of amino acids. Using a threshold of 0.5%, eight non-synonymous mutations were detected for the MPA passage (**Fig. 8B**) which included one stop codon (**Supplementary Table 2**), whereas only two non-synonymous mutations were detected for the MPA/GUO passage (**Fig. 8C**). Four of the eight non-synonymous mutations in the MPA supernatants were guanosine-replacement (C→T or G→A) mutations, whereas this was not the case for the two non-synonymous mutations in the MPA/GUO supernatants. We concluded that MPA inhibited BKPyV replication by inducing a GUO shortage that reduced supernatant viral loads and increased GUO-replacement (C→T or G→A) mutations.

**Fig. 8:**
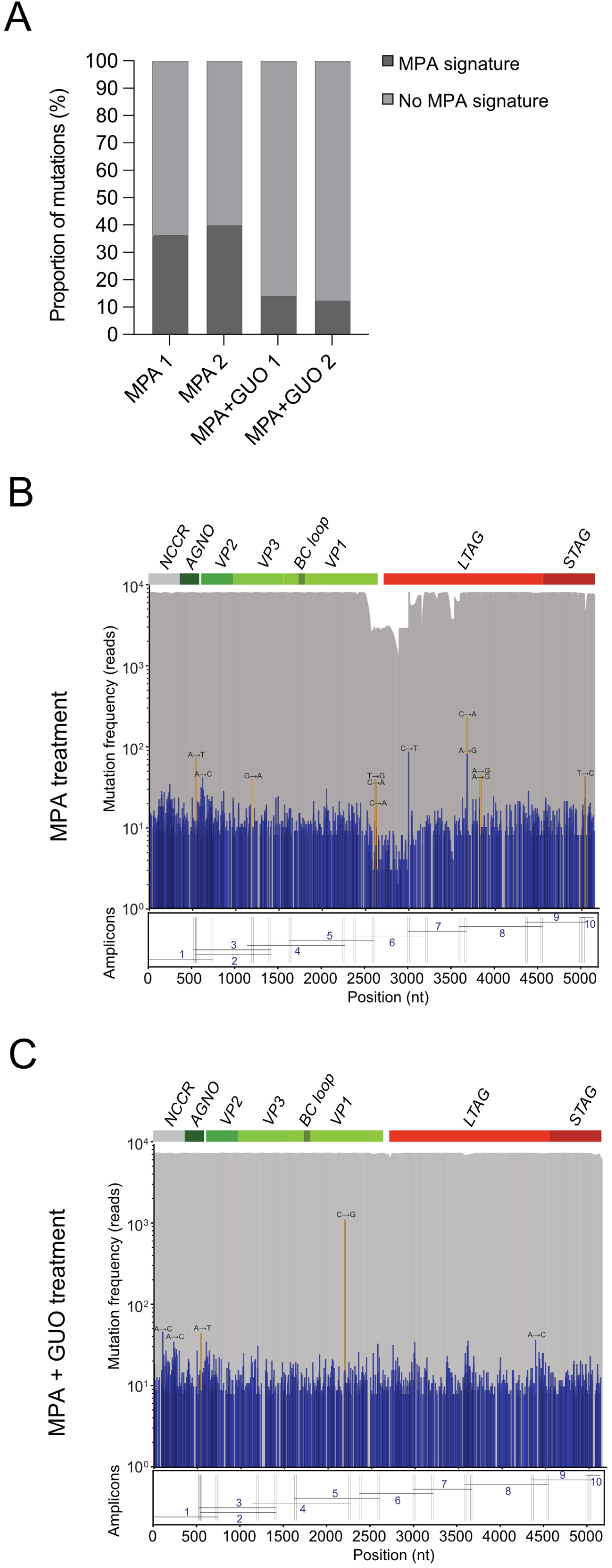
BKPyV genome mutations in supernatant viral loads.. RPTECs were seeded, infected with BKPyV and treated or with 2.5 μM of MPA (MPA) or with 2.5 μM of MPA and 200 μM of GUO (MPA/GUO) at 2 hpi. Supernatants were adjusted for BKPyV-loads and used to infect newly seeded RPTECs for three passages. Passage 3 supernatant BKPyV loads were amplified using 10 primer pairs of approximately 1000 bp covering the entire BKPyV genome. The fragments were gel-purified and submitted to next-generation sequencing (see Materials & Methods). The positions of the primers used for the 1000 bp amplification are indicated at the bottom of each diagram. NGS coverage across the BKPyV genome is shown as gray bars. Blue bars indicate the frequency of detected mutations. The orange bars represent mutations causing amino acid substitutions, shown only for variants with a frequency above 0.5%. A. Proportions of MPA signature mutations (G/C→T/A transitions; dark grey bars) and non-MPA signature mutations (light grey bars) among all detected variants in duplicate samples treated with MPA or MPA/GUO. B. BKPyV mutations identified in the supernatants of the MPA-treated RPTECs. C. BKPyV mutations identified in the supernatants of the MPA/GUO-treated RPTECs.

## Discussion

Over the past two decades, immunological risk-adapted induction and maintenance immunosuppression has significantly reduced acute rejection events in KT recipients ^11^. Thus, TAC and MPA have largely replaced cyclosporine and azathioprine as standard immunosuppression. However, the previously rare BKPyV-associated nephropathy has emerged as a regular challenge to KT outcomes ^3,40,41^, and has not been met with effective antivirals so far ^42^. Since immunosuppression is a potentially modifiable factor of BKPyV events ^24^, a better understanding of the *on*-*target* and *off-target* effects of immunosuppressive drugs may assist in optimizing current and future management strategies ^43^. In the current study, we have characterized the *off-target* effects of MPA on BKPyV-replication in primary human RPTECs that serve as a model of BKPyV-nephropathy. Our experimental study provides the following insights:

First, increasing MPA concentrations inhibit BKPyV replication with an IC_50_ of 0.65 µM for viral progeny release. MPA inhibition of viral progeny was quantified by QNAT of DNase-protected viral genomes as validated previously in experimental ^44^ and in clinical studies ^34^. Importantly, MPA-mediated reduction in DNase-resistant supernatant viral loads was associated with lowered infectious progeny that persisted over sequential cell culture passages.

Second, the MPA-mediated decline in supernatant viral loads was paralleled by a decrease in the number of BKPyV-infected cells at 72 hpi as evidenced by immunofluorescence staining of cells and by immunoblots of normalized viral and cellular proteins. In both analyses, MPA inhibition appeared to affect the *EVGR*-encoded LTag more prominently than the *LVGR*-encoded Vp1 or agnoprotein. However, the MPA dose-dependent reduction plateaued for both, the supernatant BKPyV loads as well as for the viral LTag expression, at approximately 20% of the mock-treated controls consistent with an uncompetitive inhibition pattern.

Third, the MPA-mediated decrease of LTag of 80 kDa in the immunoblots was inversely correlated with the prominent appearance of a truncTag-17. Of note, the decline in supernatant BKPyV loads, the reduction of LTag, and the appearance of truncTag-17 after MPA addition were reversed by exogenous supplementation of GUO. Finally, the reversible shortage in GUO/guanin precursors was captured by an increased rate of guanosine-replacement (C→T or G→A) mutations in supernatant BKPyV loads as compared to viral progeny in MPA/GUO-treated controls. Thus, MPA-mediated GUO depletion of primary human RPTECs can exert the partial inhibition of BKPyV replication as an *off-target* effect despite the presence of the GUO/guanin salvage pathway ^45^.

The observation that the inhibitory effect of MPA faded at approximately 20% of the solvent controls is consistent with the reported uncompetitive inhibition of the cellular IMPDH enzyme activity that is required for the *de novo* synthesis of GUO ^35,36^. This inhibition pattern precluded the determination of the MPA-IC_90_ that is predicted to mediate effective clearance of plasma BKPyV-DNAemia in KT recipients within 3 to 6 months ^43,46^. Therefore, the *off-target* effects of MPA alone are unlikely to mediate an efficient antiviral interruption of high-level BKPyV replication and clearance of BKPyV-DNAemia.

Our results also indicate that there is a subpopulation of cells that appears to be largely refractory to MPA inhibition, allowing for sustained low-level BKPyV replication and progeny release even when MPA was increased to concentrations of 4 times and more above the IC_50_ of 0.65 µM (i.e., 2.5 µM or 8 µM). We noted that the proliferation of RPTECs is also sensitive to MPA inhibition of the GUO *de novo* pathway, as shown here by microscopy and real-time impedance measurements. This observation is consistent with the fact that in proliferating cells, the *de novo* purine pathway is upregulated by the inducible IMPDH2 gene, on top of the constitutive IMPDH1 ^47^. IMPDH2 upregulation may also facilitate BKPyV replication during the cellular S-phase state, which is induced by the viral LTag promoting cellular G1→S-phase progression ^2,48–51^. Conversely, the MPA-refractory cell population consists largely of resting cells infected with BKPyV that support the sustained low-level viral replication, presumably through a limited internal turn-over feeding into the salvage pathway.

The time course studies show that MPA exposure at 24 h before infection reduced both, RPTEC confluency and supernatant BKPyV loads to a similar extent as the MPA addition at 2 hpi. With increasing delay of MPA addition after 12 hpi, the MPA inhibition faded while RPTEC confluency and supernatant BKPyV loads increased, reaching similar levels as untreated controls at 36 hpi and 48 hpi. Conversely, GUO effectively reversed the reduced supernatant BKPyV loads and cell confluency when added from −24 hpi until 12 hpi. However, insufficient GUO levels during the early phase up to the time of viral DNA replication could not be fully compensated by exogenous GUO supplementation. Thus, the levels of LTag remained reduced, whereas the truncTag-17 levels were increased supporting its role as a viral sensor of MPA-mediated GUO depletion. Interestingly, earlier work has reported the presence of several truncated variants of LTag as the result of alternative splicing, but their functional relevance including in metabolic exhaustion is awaiting further study ^52^.

Our study results are faced with several limitations. First, the MPA-inhibition of RPTEC proliferation and BKPyV replication as well as their reversal by external GUO supplementation raises the question about the relevant MPA and GUO concentrations in KT recipients. Recent studies indicate that the levels of guanine and guanosine are in the nanomolar range and therefore approximately 2- to 3-orders of magnitude below the concentrations needed to antagonize MPA-inhibition of BKPyV replication ^53^. We note that MPA is administered in KT recipients by twice daily dosing, rather than guided by trough levels used for TAC or for cyclosporin ^23,54^. However, as reviewed in detail elsewhere ^55–57^, MPA peak and trough concentrations typically range from of 7 – 37 mg/L (22 – 116 µM) and 1.2 – 3.9 mg/L (3.7 – 12.2 µM), respectively. At a rate of 3%, the free MPA concentrations likely oscillate twice daily between peak and trough concentrations of 0.66 – 3.47 µM and 0.11 – 0.37 µM, respectively. Thus, MPA exposure can reach relevant concentrations in the renal tubules via the glomerular filtrate as well as via the peritubular capillaries. These estimates are also in line with the therapeutic range discussed for MPA treatment of lupus nephritis ^58^.

Second, the results of our study are limited to a synchronized BKPyV infection model ^37,59^, whereas BKPyV-nephropathy presents as an asynchronous side-by-side of pre-, early- and late-replication phases ^48,60^. Our time course analyses suggest that approximately half of the BKPyV-replicating cells are in the late viral replication phase and no longer affected by MPA inhibition. However, MPA exposure from 24 h before until 12 h after viral infection is associated with inhibition of BKPyV replication. During these 36 hours, RPTECs are exposed to three peak concentrations in case of 12 hourly MPA administration in KT recipients. Thus, MPA is likely to affect the remaining BKPyV-replicating cells during the susceptible pre-infection and the early phase of BKPyV replication.

Third, the *off-target* effects of MPA on BKPyV replication are prominent in dividing RPTECs, on which MPA exerts a pronounced cytostatic effect. The cytostatic effect of MPA might not be relevant in people without significant tubular damage, but is likely to affect the regeneration of damaged tubular lining as encountered during advanced BKPyV pathology. When attempting to balance the undesirable anti-proliferative host cell effects and the desirable antiviral action, we believe that MPA discontinuation is appropriate for the management of KT patients with extensive viral tubular damage to permit both, the expansion of BKPyV-specific immune effectors and the regeneration of the damage renal tubule. Conversely, our data suggest that re-starting or re-increasing MPA may be well acceptable after immunologic clearance of BKPyV-DNAemia and effective renal tubule regeneration ^61^, thereby reducing the risk of subsequent rejection episodes ^3,62–64^.

The deep sequencing results highlight the need to better understand the link between immunosuppressive *off-target* effects and viral mutations in KT recipients. Recently, MPA has been reported to inhibit the SARS-CoV-2 replication, but was overcome by an increased mutation rate leading to the emergence of SARS-CoV-2 variants ^65^. Our passage experiments did not provide evidence for the emergence of MPA-resistant BKPyV variants. However, mutations of epitopes facilitating escape from T cell- or B cell-specific immunity are likely to occur, especially during prolonged high-level BKPyV replication in KT recipients treated for months and years with MPA.

We conclude that MPA inhibits both BKPyV-replication as well as its host cell proliferation, by limiting the *de novo* guanosine synthesis. This inhibitory effect can be alleviated by supra-physiologic concentrations of GUO, through the purine salvage pathway, that is constitutively absent in lymphocytes. BKPyV mutations may arise during viral replication and increase in case of GUO shortage. Our study emphasizes the importance of better integrating not only the desired *on-target* mechanisms of immunosuppressive drugs on anti-donor immunity and its unwanted side effects of equally paralyzing BKPyV-specific immune effectors, but also their *off-target* activities on non-lymphocytic and specifically virus-infected host cells ^19,30,66^. Together, our observations support the reduction or discontinuation of MPA in the current management of MPA in KT patients with new-onset BKPyV-DNAemia and nephropathy as well as re-start or re-increase of MPA after BKPyV-DNAemia clearance in the current or future management of BKPyV-nephropathy in KT recipients ^3^.

## Materials and Methods

### Cell and culture conditions

Primary human renal proximal tubular epithelial cells (RPTEC Lot:5111; 4100, ScienCell) were maintained in epithelial cell medium (EpiCM; 4101, ScienCell) and passaged with Trypsin (T3924, Sigma-Aldrich) and Defined Trypsin Inhibitor (DTI; R007100, Invitrogen) as described in detail by Manzetti et al. ^37^. Cells were passaged and re-seeded following previously published methods ^30,44^. Mycophenolic acid (MPA; M3536, Sigma-Aldrich) was dissolved in DMSO (41640, Sigma-Aldrich) at a stock concentration of 156mM. Guanosine (G6264, Sigma-Aldrich) was dissolved in DMSO at a stock concentration of 200mM.

### Virus infection

Infection with BKPyV (Dunlop strain, GenBank ID: KP412983.1) was carried out as described previously ^37^. In brief, RPTECs were seeded 24 h before infection in EpiCM (2% FBS), then cells were washed and BKPyV was diluted to a MOI of 1 in EpiCM (0% FBS) and incubated with the cells at 37°C. After 2 hours, the remaining virions contained in the supernatant were removed by washing with PBS and fresh EpiCM (0.5% FBS) was added, containing the indicated amounts of MPA or DMSO (solvent control), followed by the addition of guanosine at the indicated time points. Supernatants and cell lysates were harvested at 72 hpi for further assays.

### Quantitative nucleic acid testing (QNAT) of supernatant viral load

Total nucleic acids were extracted from supernatant according to manufacturer instructions (QIAmp Blood Mini Kit, 51104, Qiagen) after pretreatment with or without DNase-I (18047019, Thermo Fisher Scientific) to enumerate protected viral genomes as described previously ^34^. BKPyV DNA loads were quantified using a standard QNAT in triplicate in 25 µL reaction as described previously ^34,67^, using 2x qPCR Mastermix plus low ROX (RT-QP2X-03+LR, Eurogentec). QNAT was performed on the ABI 7500 HT cycler (Applied biosystems), following the program: 50°C for 2 min, 95 °C for 10 min, 95 °C for 15 sec (45 cycles) and 60 °C for 1 min (45 cycles). Results were presented as copies of genome per mL (copies/mL).

### Immunoblotting

Cells lysates were prepared in modified RIPA buffer: 10 mM Tris/HCl pH 7.5; 150 mM NaCl; 0.5 mM EDTA, 1.0% Nonidet P-40 and proteinase inhibitor (04693132001, Roche). Total protein concentrations were quantified using the BCA protein assay (23225, Pierce) and 8 µg total protein in Laemmli sample buffer (161-0747, Bio-Rad) was loaded to each well of a gradient mini gel (4-20% Mini-PROTEAN® TGX™ Precast Gel; 4561094, Bio-Rad). Gels were run in a gel electrophoresis apparatus (BioRad) at 25 mA per gel for 50 min, and proteins were then transferred onto an PVDF membrane (1704274, Bio-Rad) using the Trans-Blot Turbo Transfer System (Bio-Rad). Membranes were immersed in blocking buffer (927-40000, LI-COR, Bad Homburg, Germany) 1:2 diluted in Tris-buffered saline (TBS) for 30 min following incubation with primary (1h at RT or o/n at 4°C) and secondary antibodies (1h at RT) diluted in blocking buffer (1:2 diluted in TBS-0.1% Tween 20). The membrane was washed inbetween with TBS-0.1% Tween 20. Primary antibodies: mouse IgG2a anti-BKV SV40-LTag antibody (1:1000; ab16879, Abcam), rabbit IgG anti-Vp1 (1:5000; ab53977, Abcam), rabbit IgG anti-agnoprotein (1:1000; 1163, Eurogentec) ^37^, and mouse IgG1 anti-alpha Tubulin (1:1000; A-11126, Molecular Probes). The mouse IgG2a anti-BKV SV40-LTag antibody can detect both BKPyV LTag and truncTag, as the targeted epitope is shared between the two spliced variants. Secondary antibodies: Alexa Fluor 680 donkey anti-mouse IgG (1:15000; A10038, Invitrogen) and IRDye 800CW goat anti-rabbit IgG (1:10000; 926-32211, LI-COR). Finally, membranes were scanned in the LI-COR Odyssey imaging system (LI-COR), and images were analyzed using ImageStudioLite software (LI-COR).

Relative protein expression was determined by normalizing band intensities to the corresponding tubulin band for each condition. Where indicated, values were further normalized to the control condition, which was set to 1

### Cell proliferation

Cell proliferation of RPTECs was monitored by independent measurements using the real-time xCELLigence system (ACEA Biosciences) following the protocol described previously and by brightfield microscopy. Briefly, RPTECs were seeded in 96-well plate (xCELLigence) 24 h before infection ^38^ and the cell index normalized as indicated.

### Immunofluorescent (IF) staining

RPTECs were seeded on coverslips (Thermo Fisher Scientific) at 32500 cells/cm^2^ per 24-well plate. For improved adherence, coverslips were precoated with type I collagen (C7624, Sigma Aldrich). At time of harvest, cells were fixed with 4% paraformaldehyde at 72 hpi following the established protocol for immunofluorescent staining ^44,59^. In brief, cells were permeabilized in 0.2% Triton X-100 solution for 10 min at room temperature, and after washing, blocked with blocking buffer containing 3% bovine serum albumin (BSA) in PBS at 37°C for 15 min, following incubation with primary and secondary antibodies diluted in PBS-3% BSA. Primary antibodies: mouse IgG2a anti-SV40 LTag (1:50; clone DP02, PAb416, Calbiochem), mouse IgG1 anti-Vp1 (1:400; MAB3204-M19, Abnova), rabbit IgG anti-agnoprotein (1:750; 1163, Eurogentec) ^37^. Secondary antibodies: goat anti-mouse IgG2a Alexa 568 (1:300; A-21134 Invitrogen), goat anti-mouse IgG1 Alexa 647 (1:800; A-21240, Invitrogen), or goat anti-rabbit Alexa 488 (1:1000; ab150077, Abcam) together with Hoechst 33342 (1 µg/mL; B226, Sigma-Aldrich) for staining cell nuceli. The coverslips were then mounted on glass slides with ProLong gold antifade (P36935, Thermo Fisher Scientific), and imaged with a CELENA **®** X High Content Imaging System (20x objective).

### Supernatant virus passaging

To explore long-term effects of MPA on the viral genome, BKPyV was passaged in the presence of MPA or MPA and GUO, in duplicates. During the primary round of BKPyV infection, RPTECs were infected with stock BKPyV-Dunlop at a MOI of 1, as described above. After 72 hours, supernatants were collected from cells treated with MPA or MPA and GUO, and viral loads were quantified by QNAT. For the subsequent round of infection (passage), all input supernatants were normalized by diluting them to match the viral load of the condition with the lowest measured viral titer. Freshly thawed RPTECs were used for each passage. To monitor replication dynamics and treatment effects, this process was repeated for three passages. Viral DNA from the cell culture supernatants was extracted for PCR amplification and subsequent deep sequencing.

### Next generation sequencing of BKPyV genomes

A previously published method to generate 1000 bp amplicons was used to cover the entire BKPyV genome in 10 reactions, each consisting of 2 primer pairs (**Supplementary Table 1**)^34^. In brief, the amplification reactions consisted of 20 µL master mix (Iproof High Fidelity DNA Polymerase kit, 1725301, BioRad) containing a final concentration of 10 µM per primer pair. Five µL of extracted DNA was added per reaction and run on a Veriti™ Thermal Cycler (Applied Biosystems). Amplicons were separated on a 1% agarose gel with ethidium bromide, and bands were excised and purified using the NuceloSpin Gel and PCR Clean-up kit (740609, Macherey-Nagel). Purified amplicons were eluted in 30 µL ddH_2_O, and purified amplicons were pooled to reconstitute the complete BKPyV genome for subsequent NGS. The amplicon pools were submitted to Novogene (Cambridge, UK) for library preparation and NGS on a NovaSeq X Plus platform.

### Bioinformatics analysis

Illumina sequencing reads were processed and analyzed using a custom bioinformatics pipeline. Adapter sequences were trimmed from paired-end Illumina reads using BBDuk (tool in the BBMap package v38.98) with k-mer size of 21 and Hamming distance of 2. Trimmed reads were aligned to the reference genome KP412983 using BWA MEM (default parameters), and the resulting alignments were converted to BAM format, sorted, and indexed using SAMtools ^68^. Genome coverage was calculated using BEDTools genomecov ^69^.

The sorted BAM files were parsed directly using Pysam. At each genomic position, all reads with mapping quality >10 were extracted, and the nucleotide counts (A,T,C,G) at each position were recorded, requiring a minimum of 2 reads per variant allele and a minimum of 5 reads total. For each nucleotide position, the reference nucleotide, nucleotide frequencies, total read depth, and associated gene annotations were saved into a table. For each amino acid position, reads spanning all three nucleotide positions of the codon were identified, and codons were reconstructed, accounting for strand orientation. Mutations per position were called at a threshold of 0.5% of the position’s coverage. Database mutations were identified from the collection of sequences ^70^ and compared to the sample mutations by pairwise aligning the sequences to the reference genome KP412983. Mutations matching the APOBEC3 mutational signature were defined as C->G and C->T mutations in a TCT or TCA context.

### Statistical analysis

Data sets were analyzed with GraphPad Prism 10 software (version 10.1.0). Data sets are presented as the mean ± SD. The effective concentrations were determined using a sigmoidal 4P model, Y = Bottom + (Top-Bottom)/[1+10 (LogIC50-X) x HillSlope] in GraphPad Prism 10 software (version 10.1.0). Add statistical test used to calculate p-values.

## Acknowledgements

This work was funded in part by the personal appointment grant MM2109 to HHH from the University of Basel, Basel, Switzerland, and a SNF grant NMM1531.

Part of the results were presented as a poster at the World transplant Congress 2025, San Francisco, (Abstract# P2.05.68) and the 23^nd^ Symposium of the International Immunocompromised Host Society (ICHS) in Antalya, Turkey.

## Data availability

This data will be made publicly available as part of the article upon publication unless where selectively indicated as “data available on request” because of on-going research.

## Appendix with supporting information

Additional supporting information may be found online in the Supporting Information section.

